# Spores of the fungal pathogen *Cryptococcus* exhibit cell type-specific carbon source utilization during germination

**DOI:** 10.1101/2023.10.01.560341

**Authors:** Sébastien C. Ortiz, Megan C. McKeon, Michael R. Botts, Hunter Gage, Anna Frerichs, Christina M. Hull

## Abstract

Spores are critical morphotypes required for long-term survival of most fungi. Under the right environmental conditions, spores can escape dormancy and differentiate into vegetatively growing cells through the process of germination. For fatal human fungal pathogens like *Cryptococcus*, germination is the key differentiation process required for spores to initiate vegetative growth and ultimately cause disease; however, relatively little is known about the molecular mechanisms that control germination. To this end, we performed an extensive characterization of *Cryptococcus* spore germination through the morphological assessment of germinating spores, the inhibition of key eukaryotic processes, and the detailed quantification of fungal spore germination kinetics under numerous nutrient conditions. We identified temporal associations between molecular and morphological events and determined that carbon metabolism pathways (both glycolysis and oxidative phosphorylation) were required from the beginning of germination. We further determined that carbon sources are primarily used as fuel rather than as simply triggers of germination ‘commitment,’ and identified spore-specific carbon source utilization that is absent in yeast. Finally, we discovered the first spore-specific enzyme, Nth2, a trehalase that is required for germination when trehalose is the primary available carbon source. Together this work provides an extensive characterization of *Cryptococcus* spore germination and suggests that germination is more than simply a ‘modified cell cycle’ but is rather a highly adapted differentiation process.

## Introduction

Spores are a non-replicating, stress-resistant cell type used by many organisms to spread to new environments. To escape this seemingly dormant state, spores must sense their surroundings, establish the presence of a favorable environment, and then differentiate into vegetatively growing cells through the process of germination. Spore germination is a pivotal process because the ability to reproduce in new environments is essential to a species’ survival. The most prolific and efficient sporulators on planet Earth are fungi, and spores play essential roles in the sexual and asexual cycles of extremely diverse fungal organisms [1].

Given the importance of fungi in nearly all ecosystems, and the critical role that spore germination plays in fungal survival, it is surprising how little is known about the molecular mechanisms that control the initiation and maintenance of germination. This gap is particularly acute for the germination of human fungal pathogens, the majority of which initiate infections through the inhalation of spores [2–4]. For these pathogens, germination into vegetatively growing cells is required for spore-mediated disease, yet limited studies have been carried out to understand this key process. One possible explanation for limited inquiry into spore germination stems from studies suggesting that germination is simply a modified cell cycle designed to escape any relatively static growth state (e.g. stationary phase growth) [5,6]. This impression, largely from studies of *Saccharomyces cerevisiae* spores, may have deterred molecular biologists from viewing germination as an area of biology with new opportunities for discovery.

Recent evidence from the human fungal pathogen *Cryptococcus*, however, suggests that spore germination is a specific process with properties and components distinct from the canonical cell cycle. In particular, a proteomic analysis comparing spores and yeast of *Cryptococcus* identified 18 spore-enriched proteins, 7 of which had no identifiable homology to any characterized proteins [6]. When the gene encoding one of these proteins, Identified <u>S</u>pore <u>P</u>rotein 2 (Isp2), was deleted, the resulting strains produced spores that demonstrated a unique germination defect, suggesting that spore germination (at least in *Cryptococcus*) is not simply a modified cell cycle but is rather a fungal-specific differentiation process that has yet to be characterized. As such, germination could involve undiscovered molecules and pathways in eukaryotic biology and provide novel fungal-specific targets ideal for the development of new antifungal therapeutics.

Another reason that germination is poorly characterized overall may be a general lack of tools available to fully probe spore biology. Recent efforts have been made to develop *Cryptococcus* as a model for studying fungal spore germination [8, 9]. The *Cryptococcus* system has several advantages, including a method for bulk spore purification and a high-resolution quantitative germination assay (QGA) that allows independent assessments of germination kinetics of thousands of spores simultaneously. Using this assay, population-level germination phenotypes that correlate with specific molecular pathways have been identified, which adds to the extensive molecular and genetic tools of the *Cryptococcus* system [9].

To determine key properties and events required for successful germination of fungal spores, we took advantage of these advanced tools to carry out an evaluation of *Cryptococcus* spore germination. We performed a morphological assessment of germinating spores, probed gemination with a variety of inhibitors of key eukaryotic processes and carried out a detailed characterization of fungal spore germination kinetics under numerous nutrient conditions. We identified spore-specific metabolic activity that is absent in yeast and discovered the first spore-specific enzyme, Nth2, a trehalase that is required for germination when trehalose is the primary available carbon source. This characterization of germination initiation and kinetics provides critical insights into the fundamental events required for spores to differentiate, survive, and ultimately establish vegetative growth in new environments.

## Results

### Spore germination encompasses changes in cell morphology, intracellular architecture, and surface carbohydrate composition

To determine physical changes in spores over the course of germination, four microscopy-based approaches were used to characterize populations of germinating spores (Figure 1).

**Figure 1.**
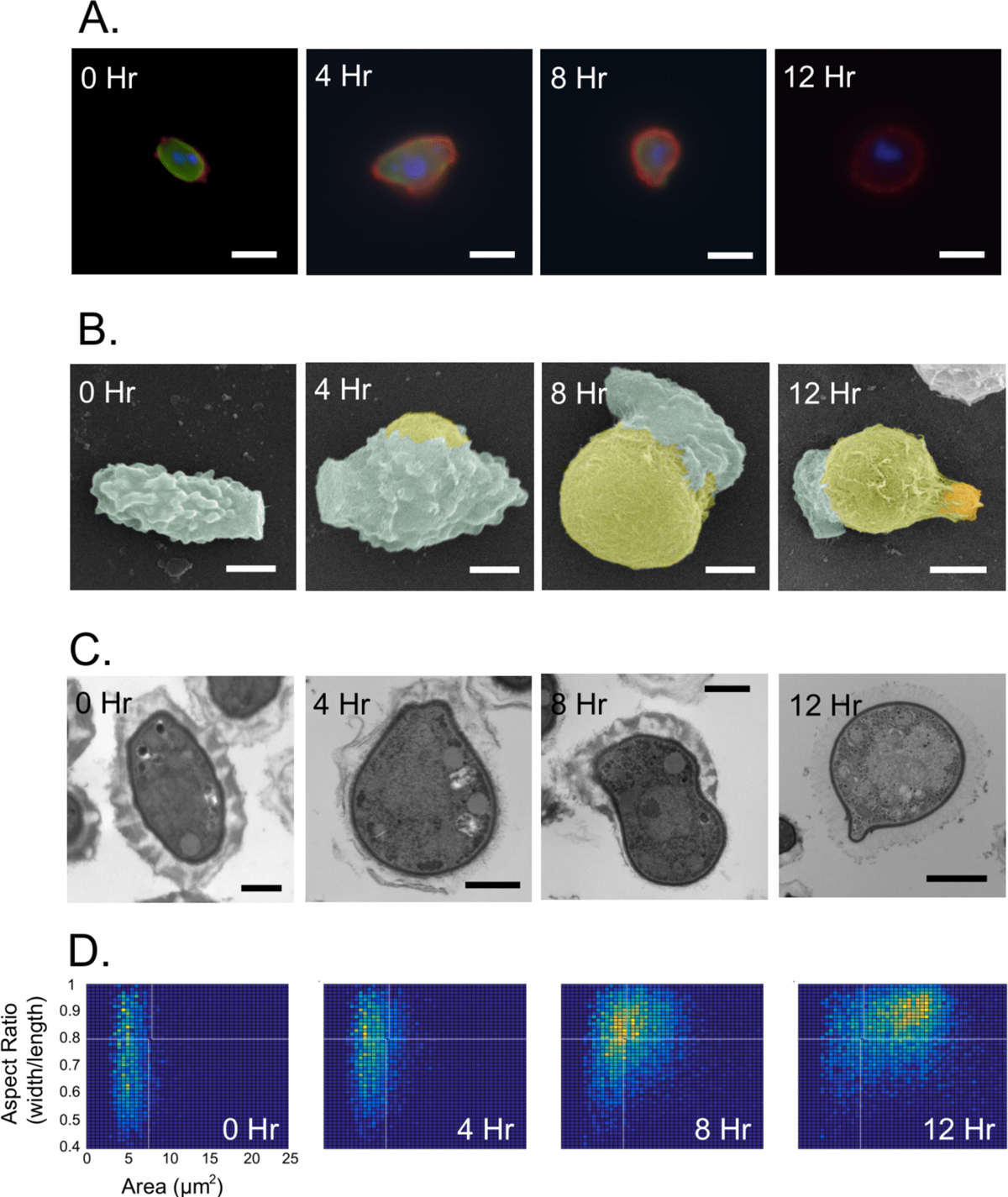
Microscopic evaluation of spore germination demonstrates reproducible changes in spore size, shape, and surface properties. Images of spores at 0, 4, 8 and 12 hours after addition of 2% (111 mM) glucose. A) Representative fluorescent images of germinated spores fixed and stained with with DAPI (blue), *Datura stramonium* lectin (green) and anti-glucoronoxylomannan antibody (red). White bars represent 0.75 µm. B) Representative false-colored scanning electron microscopy (SEM) images of germinating spores (mint), showing emergence of a yeast (yellow) from the side of the spore to produce a yeast with an attached spore remnant that buds off daughter cells (orange). White bars represent 1 µm. C) Representative images of transmission electron microscopy (TEM) of germinating spores demonstrating a transition from small, oval cells with a thick surface layer to a larger, round yeast with a canonical capsular surface. Black bars represent 0.75, 1, 2, and 2.5 µm across panels from left to right. D) Two dimensional histograms from quantitative germination assays (QGA) showing population-level morphological changes of ∼6000 germinating spores. The x-axes represent size from 0 to 25 μm^2^; the y-axes represent aspect ratio (width/length) from 0.4 to 1.

Previous studies of spores and yeast identified cell type-specific exposed carbohydrates that we hypothesized would change in a reproducible pattern as spores transitioned into yeast [10,11]. Using probes to cell type-specific markers, we monitored changes in surface epitopes over the course of germination at 0, 4, 8, and 12 hours in rich culture medium (YPD). At each time point, cells were fixed with ethanol and stained with 1) an *N*-acetylglucosamine-binding lectin (from *Datura stramonium*) tagged with fluorescein (DSL-green), 2) a monoclonal antibody to glucuronoxylomannan (GXM) tagged with rhodamine (anti-GXM-red), 3) and the DNA binding molecule 4′,6-diamidino-2-phenylindole (DAPI) to identify nuclei (blue). Using fluorescence microscopy, we observed that spores consistently fluoresced green homogeneously across the cell surface at time 0 and gradually lost the lectin signal over the 4, 8, and 12 hour time points. By 12 hours, there was no visible DSL-green signal, indicating no discernable DSL-bound *N*-acetylglucosamine on the cell surface. In contrast, we observed that the highly punctate pattern of anti-GXM antibody binding to spores gradually transitioned to homogeneous binding across the germinated spore surface over time, indicating that the distribution and/or amount of GXM changed as spores transitioned into yeast (Figure 1A). DAPI staining indicated that the vast majority of spores harbored a single blue punctum, consistent with mononucleate spores. However, spores occasionally showed a secondary blue punctum, suggesting the presence of a second nucleus (Figure 1A, Time 0 Hr). To determine the number of spores with more than nucleus, fluorescently stained nuclei were scored visually in 164 spores; 157 showed a single punctum and were designated mononucleate (95.7%), and 7 showed two puncta and were designated as likely binucleate (4.3%). These percentages are consistent with prior reports that a low percentage of spore progeny from microdissections germinate to produce diploid yeast [12]. To assess changes in cellular morphology during germination, electron microscopy was carried out on germinating spores. Scanning electron microscopy (SEM) revealed crenulated, rough spores germinating by bulging consistently from one longitudinal side to ultimately produce a yeast (with a spore remnant attached). The resulting founder yeast then produced a bud perpendicular to the spore remnant (Figure 1B). Transmission electron microscopy (TEM) revealed a transition from small, oval spores with a thick spore coat composed of variable electron-dense and electron-light regions, to a larger, round yeast covered in a previously described fibrillar-appearing capsule layer of relatively low electron density (Figure 1C). To determine whether the emerging yeast were formed using existing chitin from spores or from new chitin synthesis, we stained spores with calcofluor white, washed, and imaged them over the course of germination (Supplemental Video 1.1, 1.2). We anticipated that newly produced chitin (which had not been present during calcofluor white staining) would not fluoresce, whereas chitin associated with the spore coat would remain fluorescent throughout germination. During germination we observed that visible calcofluor white staining was largely lost at the site of yeast emergence along the longitudinal sides of germinating spores and was retained at the site of the spore remnant. The protruding yeast harbored little visible fluorescence, suggesting that the yeast cell wall was formed primarily by newly synthesized chitin. Additionally, we observed that daughter cell budding from spore-derived yeast occurred repeatedly from the same site, suggesting a unipolar budding pattern (Supplemental Video 1.1, 1.2). This is distinct from the budding pattern of haploid yeast, which show bipolar budding, as demonstrated by bud scar staining and axial budding shown (Supplemental Figure 1). This unique budding pattern suggests that early spore-derived yeast and vegetatively growing yeast could be distinct cell types and may exhibit different biology.

Finally, we used a quantitative germination assay (QGA) to evaluate morphological changes across a population of germinating spores. A population of ∼6,000 spores was exposed to rich growth medium (SD) and measured for area and aspect ratio every four hours. The resulting data were plotted in two-dimensional histograms and show that spores as a population at time 0 were relatively small and oval. At 4 hours of germination, the population of spores became rounder, and at 8 hours of germination the spore population became both rounder and larger. At 12 hours of germination, the spore population had transitioned fully into round, large yeast, demonstrating a synchronous increase in circularity and size over the course of germination (Figure 1D). These observations of morphological and compositional changes provide a general understanding of the changes that occur during differentiation of spores into yeast on both single-cell and population-based levels.

### Inhibition of key eukaryotic processes during germination results in process-specific phenotypes

To evaluate the roles of fundamental eukaryotic processes during germination, inhibitors of those processes were evaluated for their effects on germination over time. It has been shown previously that the eukaryotic ribosome inhibitor cycloheximide, as well and other translation inhibitors (i.e., geneticin – G418, anisomycin, and puromycin), are potent inhibitors of *Cryptococcus* germination [9]. Inhibition of translation causes an overall “slow down” immediately from the onset of germination in quantitative germination assays, suggesting that translation is required very early in the germination process [9]. We hypothesized that chemical inhibition of other molecular events important for germination would also cause changes in population-level behaviors. To identify other fundamental processes required for germination, we used the same approach to test inhibitors of glycolysis (2-deoxyglucose), transcription (actinomycin D), splicing (pladienolide B), histone acetylation (CPTH2), DNA replication (aphidicolin), and histone deacetylation (trichostatin A). In all cases spores were grown in synthetic dextrose (S + 2% D-glucose) medium in the presence of a range of concentrations of each inhibitor and evaluated using QGAs.

Spores exposed to the non-metabolizable glucose analog 2-deoxyglucose (2-DG) at 125 mM and higher showed inhibition of germination, demonstrating an asynchrony phenotype in which individual spores germinated at different efficiencies (Figure 2A). This inhibition was observed in a concentration-dependent manner with full inhibition observed at 500 mM 2-DG. This result suggests that glycolysis is essential from the start of germination. It has been shown previously that an inhibitor of complex III of the electron transport chain, antimycin A, also causes an asynchrony phenotype at low concentrations [9]. At higher concentrations (80 µM) antimycin A, like 2-DG, fully inhibited germination. Together, these data indicate that key pathways in glucose metabolism (glycolysis and oxidative phosphorylation) are important in germination from at least as early as the beginning of the morphological transition out of the spore state (occurring between 2 and 4 hours after exposure to glucose).

**Figure 2.**
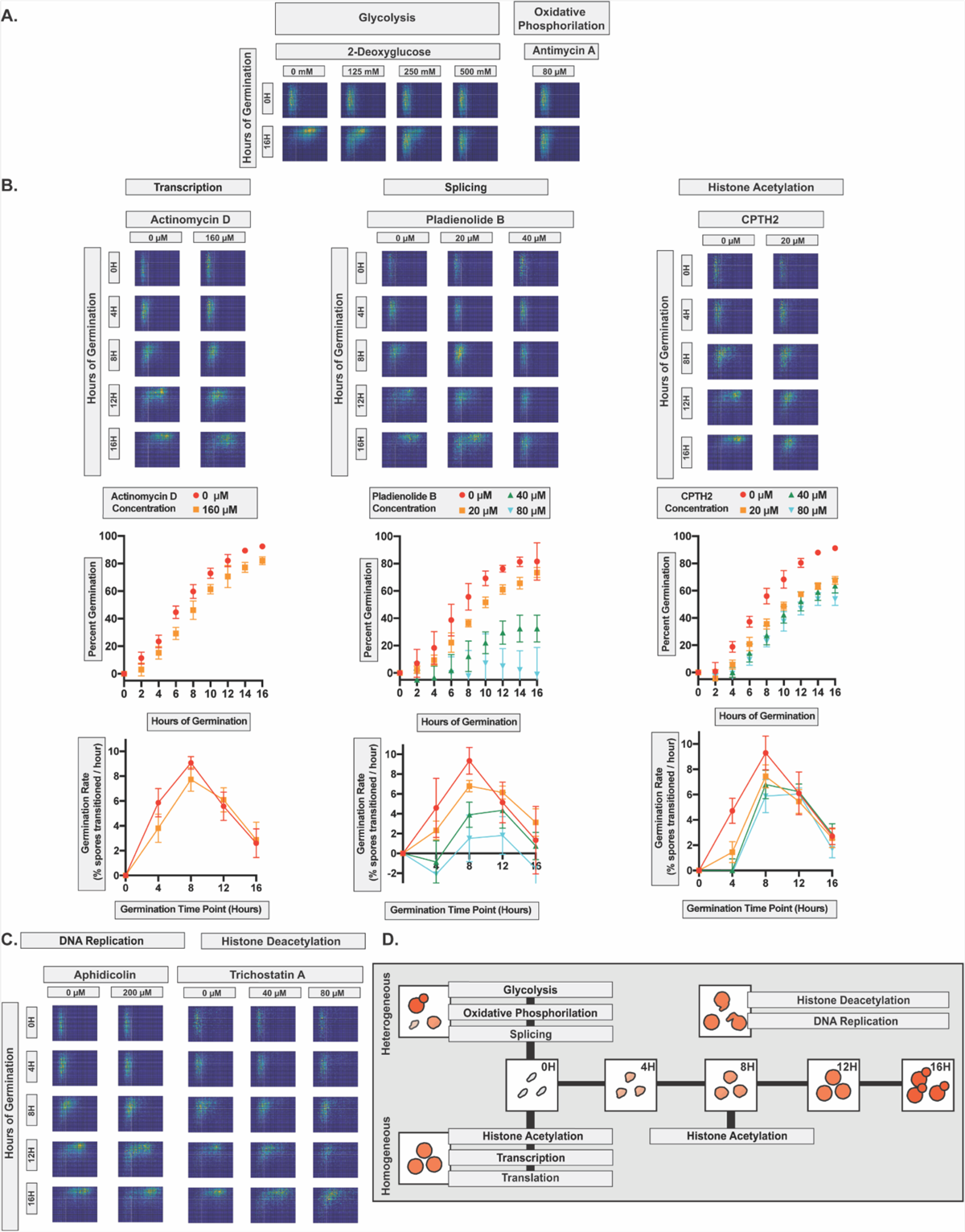
Inhibition of key eukaryotic processes during spore germination results in discernable population-level phenotypes. Evaluation of the effects of germination inhibitors on ∼6000 spores/assay using the QGA. **A)** Germination profiles of spores at 0 and 16 hours of germination in the presence of 2-deoxyglucose (125, 250, and 500 mM) or antimycin A (80 μM). **B)** Germination profiles of spores at 0, 4, 8, 12 and 16 hours of germination in the presence of actinomycin D (160 μM), pladienolide B (20 and 40 μM), or CPTH2 (20 μM), along with quantitative assessments of germination and germination rates for actinomycin D (160 μM), pladienolide B (20, 40 and 80 μM), and CPTH2 (20, 40 and 80 μM). **C)** Germination profiles of spores at 0, 4, 8, 12 and 16 hours of germination in the presence of aphidicolin (200 μM) or trichostatin A (40 and 80 μM). **D)** Diagram of the temporal roles of each probed eukaryotic process during germination. For all two-dimensional histograms, the x-axis represents cell size from 0 to 25 μm^2^, and the y-axis represents aspect ratio (width/length) from 0.4 to 1.

We also discovered that when transcription was inhibited by actinomycin D at 160 µM, germination was slowed from the start but remained synchronous across the population. These results demonstrate that, like new protein synthesis, new transcription appears important very early in germination (Figure 2B). Similarly, when splicing was inhibited by the SF3B1-specific compound pladienolide B at 20 µM, germination became asynchronous, and at higher concentrations (over 40 µM), germination was fully inhibited. This suggests that splicing is an essential process during germination that also occurs very early.

When histone acetylation was inhibited by CPTH2 at 20 µM and higher, a unique phenotype was observed in which germination was initially slowed, resumed at a normal rate, and then stalled as small circular cells (equivalent to spores after 8-10 hours of efficient germination). CPTH2 was unable to inhibit germination fully, showing identical germination kinetics at 20, 40, and 80 µM. These results suggest that histone acetylation promotes efficient initiation of germination and plays a critical role in the isotropic growth phase of germination. When histone deacetylation was inhibited by trichostatin A at 40 µM (Figure 2C), germination proceeded normally until ∼8 hours of germination. After 8 hours, spores were able to undergo isotropic growth, but in a non-synchronous manner, leading to morphological variability in the population. This isotropic growth was inhibited fully at 80 µM.

The inhibition of DNA replication by aphidicolin at 200 µM led to a phenotype similar to trichostatin A in which germination occurred normally in the first 8 hours followed by morphologically variable isotropic growth. This finding suggests that DNA replication is important during the later phase of germination when small, circular cells become larger. Other inhibitors of eukaryotic processes were tested and did not inhibit germination under the concentrations and conditions tested. These included the proteasome inhibitor bortezomib (650µM), the DNA methylase inhibitors 5-azacytidine (50µM) and sodium butyrate (80µM), the DNA replication inhibitor hydroxyurea (80µM), the microtubule inhibitors nocodazole (80µM) and paclitaxel (80µM), and the actin polymerization inhibitor cytochalasin A (80µM) (Supplemental Figure 2). These data suggest that the processes inhibited by these compounds are not required for germination; however, we cannot rule out the possibilities that these inhibitors are unable to enter the spores to cause inhibition or do not inhibit at the concentrations tested.

Overall, these results indicate that the key cellular processes of glycolysis, oxidative phosphorylation, transcription, splicing, translation, histone acetylation, histone deacetylation and DNA replication are all required for efficient spore germination to occur. Furthermore, the importance of any given process is manifested at distinct morphological stages and time points, resulting in discernable population-level phenotypes, likely indicative of the time during germination at which each process is first required to act.

### Carbon limitation leads to synchronous and concentration-dependent stalling of spore germination

Previous experiments established that germination proceeded efficiently in the presence of glucose and a nitrogen source at 30 °C [8]. However, the effects of other metabolic and environmental conditions on initiation and maintenance of germination were unknown. To establish a baseline germination response under controlled nutrient conditions, we assessed spores at 30°C in the presence of a buffered solution (1x PBS alone), S medium (Synthetic medium without glucose), glucose alone (111 mM in 1x PBS) and in SD medium (Synthetic medium with 111 mM glucose) (Figure 3A). Spores exposed to 1x PBS alone were unable to germinate after 16 hours as demonstrated by a complete lack of transition out of the spore state. In the presence of S medium, which contains 5 mg/mL ammonium sulfate and trace metals, spores demonstrated a slight increase in circularity after 16 hours but did not show efficient germination. In the presence of glucose alone, spore germination was initiated and maintained over 16 hours, but at a slower rate than in nutrient-replete conditions (SD). Together, these results demonstrate that both nitrogen and glucose are important for efficient initiation and maintenance of germination, and the presence of glucose as a carbon source plays the larger role in the efficiency of this transition.

**Figure 3.**
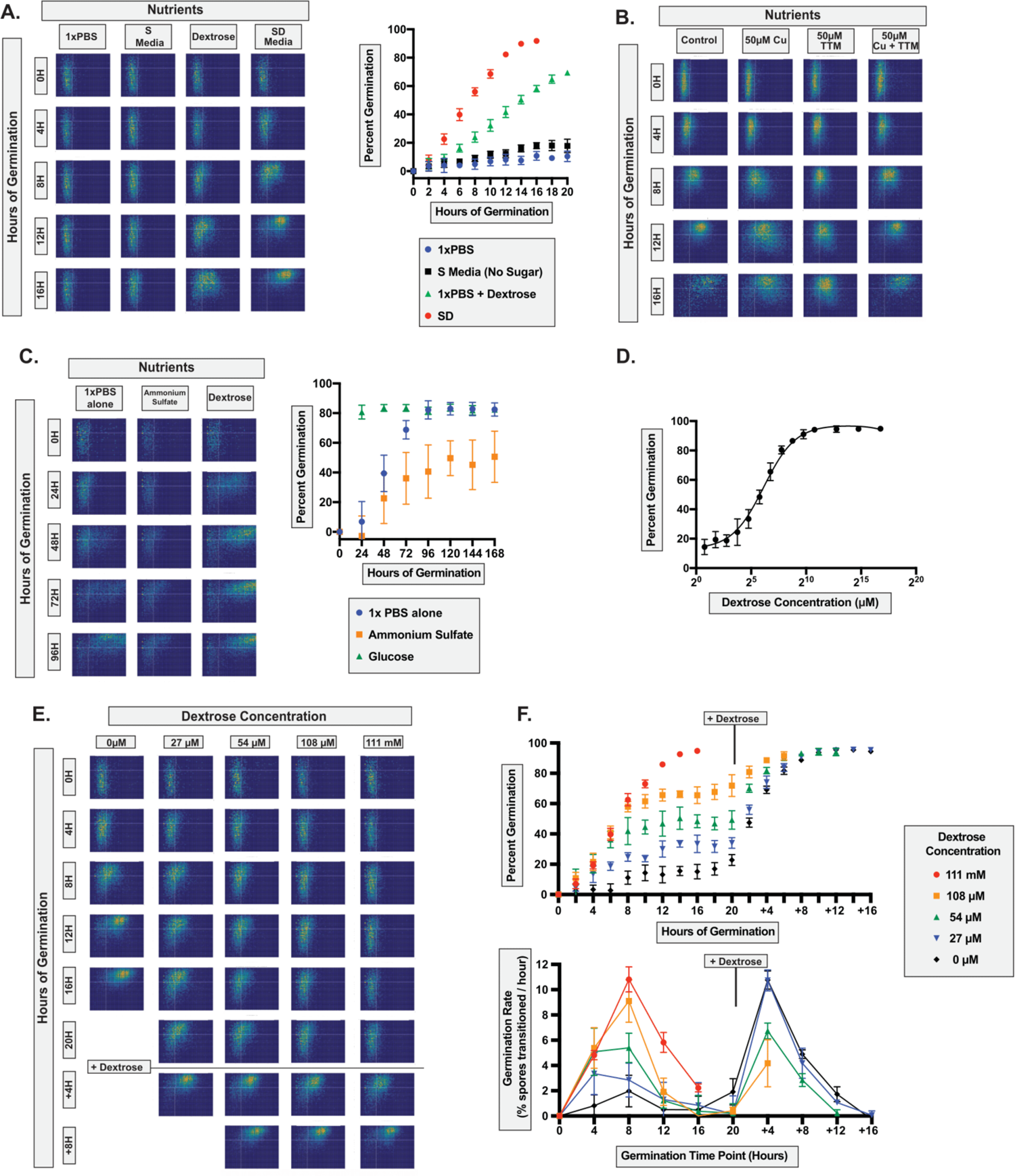
Carbon source limitation leads to synchronous and concentration-dependent stalling of germination. Evaluation of germination under a variety of nutrient limiting conditions using the QGA. **A)** Germination profiles of spores at 0 and 16 hours of germination in 1X PBS, S medium (no carbon source), glucose (111mM), or SD (left) and a graph of percent germination vs. time (right). **B)** Germination profiles of spores at 0, 4, 8, 12, 16 hours of germination in SD alone or supplemented with excess copper (50 μM CuSO4), a copper chelator (50 μM TTM) or both. **C)** Germination profiles of spores evaluated every 24 hours for seven days in the presence of 1X PBS alone, ammonium sulfate (5 mg/mL), or glucose (111mM) (left) and a graph of percent germination vs. time (right). **D)** Dose response curve evaluating the efficiency of germination at 16 hours of germination in 1.7 μM to 111mM glucose in SD medium. **E)** Germination profiles of spores at 0, 4, 8, 12, 16, and 20 hours of germination under glucose limiting conditions (27 μM, 54 μM, and 108 μM) followed by addition of excess glucose. **F)** Quantification of germination stalling and re-initiation, quantifying both relative germination (top) and germination rates (bottom). For all two-dimensional histograms, the x-axis represents cell size from 0 to 25 μnm^2^, and the y-axis represents aspect ratio (width/length) from 0.4 to 1. In all graphs Percent Germination indicates the percentage of spores that have escaped the spore state. Germination Rate indicates percent escape from the spore state per hour over time.

A metal that has been heavily implicated in many aspects of *Cryptococcus* biology, including sexual development and virulence, has been copper [13, 14]. To assess the role of metals in germination, copper (50 μM CuSO_4_) was supplemented to SD medium, and germination was evaluated (Figure 3B). Germination was efficient under these conditions but showed a higher level of morphological variability (increased variations in sizes and shapes). To evaluate copper limiting conditions, the highly specific copper chelator tetrathiomolybdate (TTM) was tested at 50 μM and resulted in efficient germination initially but stalled at the isotropic growth phase of germination. Upon addition of both 50 μM TTM and 50 μM CuSO4, germination remained fully efficient with population behavior comparable to the control. Other non-specific metal chelators, DIP and 1,10-phenanthroline, showed similar phenotypes, with 1,10-phenanthroline demonstrating a clear concentration dependent stall of germination (Figure S3). Together these results suggest that metals play a key role in germination and that metal limitation can lead to stalling of germination.

Previous reports have suggested that *Cryptococcus* spores can germinate efficiently in the apparent absence of nutrients [15], which contrasts with our findings that spores did not germinate in 1x PBS after 16 hours at 30°C (Figure 3A). To further characterize the nature of spore germination in the absence of nutrients, we evaluated germination in the presence of a buffered solution (1x PBS alone), 5 mg/mL ammonium sulfate, or 111 mM glucose every 24 hours for 7 days at 30°C (Figure 3C). We discovered that in the presence of PBS alone, spores could germinate slowly with low efficiency, low synchrony, and more morphological variability than under nutrient rich conditions. By 96 hours of incubation most spores were fully germinated, and the resulting population of yeast showed substantial size variability.

In the presence of ammonium sulfate, initiation of spore germination remained synchronous, but spores were unable to grow isotopically, suggesting that the presence of nitrogen alone is insufficient to sustain germination and facilitate replication. Finally, in the presence of glucose alone, most of the population was able to germinate fully in a synchronous manner. In each of these conditions, a small proportion of spores was unable to germinate, supporting the hypothesis that there is variation in germination ability across spores when they are in nutrient limiting conditions. Together, these data demonstrate that nitrogen, metals, and carbon are all essential for efficient germination; however, in the absence of nutrients, spores are capable of spontaneous germination that is both inefficient and asynchronous.

Based on these data, it appears that although nitrogen and metals provide optimal conditions for germination, carbon source utilization is the primary regulator of efficient germination initiation and maintenance. In other systems, brief exposure to an activating nutrient such as glucose (in the case of *Saccharomyces cerevisiae*) or L-alanine (in the case of *Bacillus subtilis*), is sufficient to induce germination, which is maintained even after the activating nutrient is removed. This signal-based induction of germination has been termed “commitment” [16–18]. To evaluate germination commitment in *Cryptococcus*, spores were exposed to a broad range of glucose concentrations from 1.7 µM to 111 mM (in synthetic medium). We determined that spore germination occurred efficiently at glucose concentrations at and above 434 µM. At lower concentrations (i.e., glucose limiting conditions) germination was hindered in a concentration dependent manner (Figure 3D). In glucose limiting conditions, spores initiated germination with normal efficiency but then abruptly stopped germinating in a concentration-dependent manner in which lower concentrations of glucose resulted in earlier stalls in germination (Figure 3E, F). These data suggested that glucose availability was required to sustain germination. To test this hypothesis, we carried out germination assays for 20 hours in limiting glucose and then supplemented with excess glucose (111 mM). Spores that had stalled during germination in concentrations of glucose below 434 µM resumed efficient germination at a normal rate, regardless of their germination state/morphology. These results suggest that spores use glucose as a source of fuel and that when glucose runs out, spore populations stop germinating. Upon addition of more fuel, the germination process can proceed. Together, these data suggest that carbon sources (at least glucose) are used as fuels during *Cryptococcus* germination as opposed to acting only as signals to induce germination and that carbon sources may need to be imported and metabolized to induce a morphological change.

### Spores can metabolize carbon sources for germination that yeast cannot metabolize for vegetative growth

It is well established that the ability of organisms to break down and metabolize different carbon sources varies significantly across species. Spores are necessary for fungal survival and likely need to be metabolically versatile to survive in new environments. We therefore hypothesized that spores would sense, breakdown, import, and metabolize a wide variety of carbon sources. To evaluate the metabolic potential of spores, we monitored the germination kinetics of spores in the QGA in the presence of different carbon sources and evaluated yeast metabolic potential in tandem using standard liquid growth assays (Figure 4, Figure S4, Figure S5, and Table S1).

**Figure 4.**
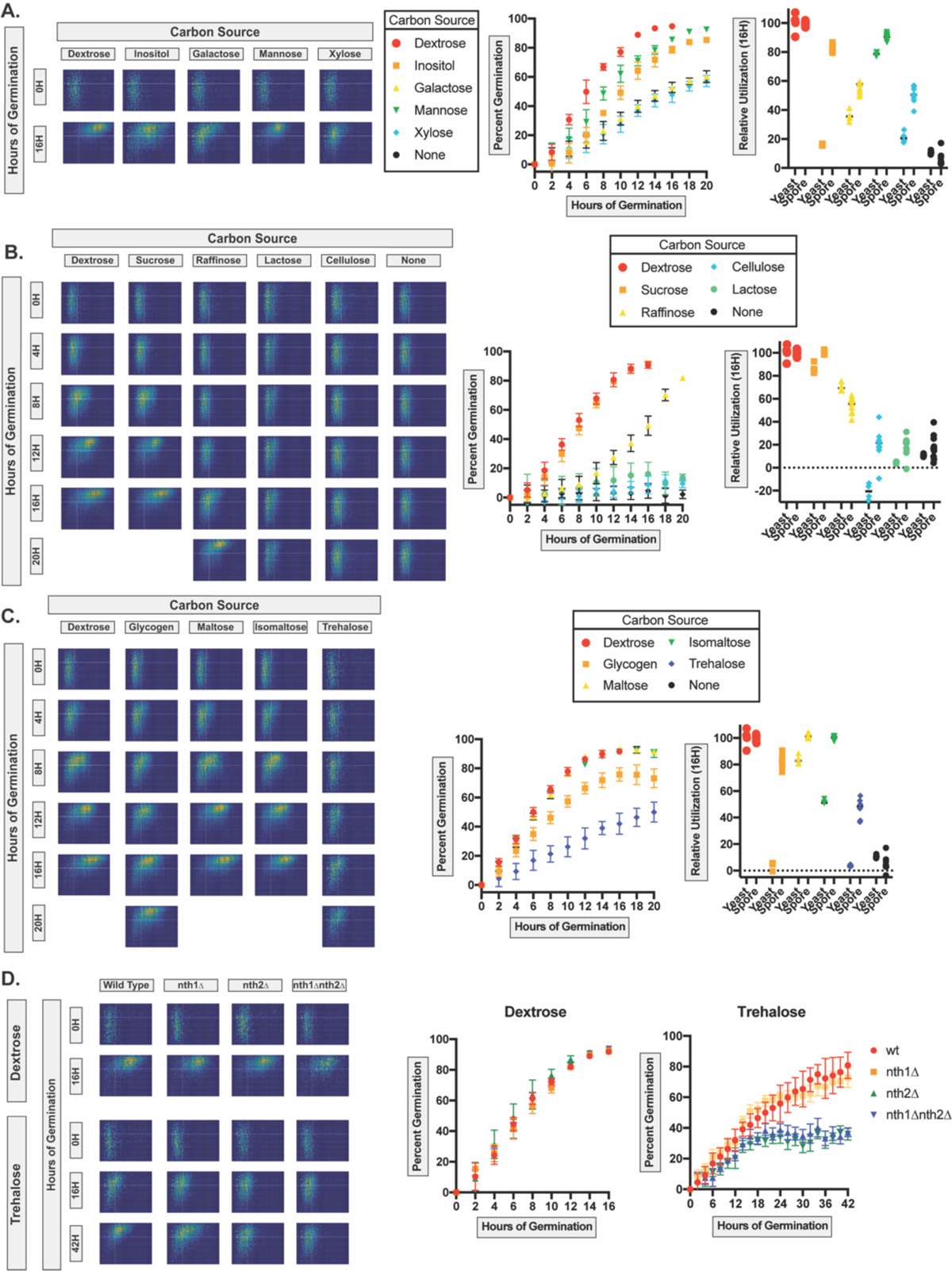
Spores demonstrate a different metabolic potential than yeast, resulting in spore-specific carbon source metabolism. Evaluation of germination in the presence of a variety of sugars as sole carbon sources (at 111 mM) using the QGA. **A)** Germination profiles of spores in the presence of simple sugars with known roles in *Cryptococcus* biology: inositol, galactose, mannose, and xylose at 0 and 16 hours (left), graph of germination over time (0 to 20 hours) (middle), and plot comparing with yeast growth (right). **B)** Germination profiles of spores germinating in the presence of complex sugars: sucrose, raffinose, lactose and cellulose at 0, 4, 8, 12, 16 and 20 hours of germination (left), graph of germination over time (middle), and plot comparing with yeast growth (right). **C)** Germination profiles of spores germinating in the presence of the complex sugars: glycogen, maltose, isomaltose, and trehalose at 0, 4, 8, 12, 16 and 20 hours of germination (left), graph of germination over time (middle), and plot comparing with yeast growth (right). **D)** Germination profiles of wild type, *nth1Δ*, *nth2Δ*, and *nth1Δnth2Δ* spores germinating in the presence of glucose (at 0 and 16 hours - top of left panel) or trehalose (at 0, 16 and 42 hours - bottom of left panel) as the sole carbon sources, graphs of germination over time in glucose (middle), or trehalose (right) as the sole carbon sources. For all two-dimensional histograms, the x-axis represents cell size from 0 to 25 μm^2^, and the y-axis represents aspect ratio (width/length) from 0.4 to 1. In all graphs Percent Germination indicates the percentage of spores that have escaped the spore state.

In addition to glucose, a handful of simple sugars have been implicated as important in the *Cryptococcus* life cycle. One of these is inositol, which has been reported to play a role in sexual development and virulence. *Cryptococcus* is known to have 10-11 inositol transporters, further indicating the importance of this sugar in its biology [19–21]. Three other simple sugars are biologically relevant to *Cryptococcus*, including galactose, xylose, and mannose, which are components of the yeast capsule (GXM and GalXM) and important to yeast biology and virulence [22]. Each of these sugars was tested as the sole carbon source at 111 mM in S medium for its ability to sustain either spore germination or yeast growth (Figure 4A). In the presence of inositol, spores germinated at a slightly slower rate than in glucose and with more morphological variation. Spores germinated in inositol showed 83.0% germination relative to spores in glucose after 16 hours, suggesting that inositol is a relatively good carbon source for germination. When inositol was assessed as the sole carbon source for yeast replication however, yeast were unable to replicate efficiently (as determined by OD_600_) showing only 16.0% yeast growth relative to glucose after 16 hours. These results suggest that spores have an ability to utilize inositol as a carbon source that is not active in their yeast parents (Table S1).

In contrast, in galactose, spores did not germinate well or grow efficiently, demonstrating 55.7% relative germination and 36.1% relative yeast growth when compared to the glucose control. Additionally, spores in galactose germinated asynchronously, suggesting that spores within a population vary in their abilities to import and/or metabolize galactose. In mannose, spores germinated efficiently (90.4% relative germination), and remained synchronous throughout germination, suggesting that mannose is a readily transported and metabolizable carbon source. Yeast showed 78.5% growth in mannose relative to glucose. In the presence of xylose as the only carbon source, spores did not germinate efficiently (50.4% relative germination), but unlike galactose-induced germination, spores in xylose remained synchronous, indicating that xylose is not used efficiently but is accessed similarly by all spores in the population. Yeast showed 21.2% growth in xylose relative to glucose. Together, these data indicate that spores harbor the ability to germinate with different levels of efficiency in response to many simple sugars known to play key roles in *Cryptococcus* biology. Additionally, germination in different sugars indicates that not all spores in a population respond equally to all sugars, leading to asynchronous germination in some cases. Finally, spores appear more efficient at utilizing these sugars for germination than yeast do for growth, particularly in the case of inositol. Five additional simple sugars (fructose, L-arabinose, D-arabinose, ribose, and rhamnose) were tested for their abilities to support germination and vegetative growth (Figure S4A). Fructose induced germination with an efficiency equivalent to glucose and was indistinguishable from glucose as a carbon source for yeast growth. In contrast, L-arabinose, D-arabinose, ribose, and rhamnose were more efficient at sustaining germination than at supporting yeast growth, showing at least a 20% difference in efficiency between germination and growth. Five non-sugar carbon sources (acetate, pyruvate, succinate, glycerol, and fumarate) were all tested for their abilities to induce and maintain efficient germination (Figure S4B). Pyruvate, succinate, and fumarate were unable to induce germination or sustain yeast replication. Acetate was able to induce germination, but it was inefficient (22.5% germination relative to glucose) and sustained yeast growth at a similar (low) efficiency (18.2% relative growth). Spores used glycerol at an efficiency similar to acetate (26.2% relative germination); however, yeast were largely unable to utilize glycerol for growth (6.7% relative growth). Together these results demonstrate variable effects of different carbon sources on germination, suggest that some carbon sources are more readily metabolized than others, and indicate that spore carbon source utilization is distinct from yeast.

To test the abilities of spores and yeast to metabolize more complex sugars, we evaluated the germination efficiency of spores in the presence of four additional carbohydrates: sucrose, raffinose, cellulose, and lactose (Figure 4B). Sucrose is a disaccharide of glucose and fructose, the two most efficient monosaccharides at inducing/maintaining spore germination. Sucrose was able to induce and maintain germination as efficiently as both glucose and fructose (101.1% relative germination efficiency), suggesting that spores harbor surface-exposed or secreted invertase enzymes that readily break α-1, β-2-glycosidic bonds. Yeast also demonstrated relatively high utilization of sucrose (86.6% relative yeast growth). Raffinose is a trisaccharide of glucose, fructose, and galactose, (effectively a sucrose molecule with a 1,6-α bond to galactose). To access raffinose as a carbon source, organisms generally must produce an α-galactosidase to liberate galactose before the glucose-fructose bond can be cleaved [23]. In the presence of raffinose as the sole carbon source, we found that visible germination was not initiated for the first 6-8 hours. At ∼8 hours after exposure to raffinose, spores began to initiate germination at the same rate as spores in glucose. Thus, until ∼8 hours after exposure to raffinose the population of spores demonstrated a morphology equivalent to 0 hours in glucose. Fourteen hours in raffinose appeared equivalent to ∼6-hours in glucose, and by 20 hours in raffinose, spores had germinated to the equivalent of spores after ∼12-hours in glucose. This result suggests that spores cannot immediately access raffinose as a carbon source but acquire the ability to do so ∼8 hours after initial exposure. One possibility for this phenotype is that spores require 6-8 hours to produce or access the required α-galactosidase and/or transporters to metabolize raffinose into galactose and sucrose. In raffinose, yeast demonstrated a similar delay in replication; however, the delay was shorter, taking only 4-5 hours to initiate growth (Figure S5). Both cellulose and lactose were unable to induce germination or yeast growth, suggesting that they cannot be imported and/or metabolized by either spores or yeast.

Two complex sugars composed of glucose subunits, glycogen (composed of maltose and isomaltose) and trehalose, are known to be used as storage molecules in other fungi [24,25]. Due to their unique roles in storage and their implications in spore biology, we hypothesized that spores would be able to readily metabolize these storage molecules [24,25]. To test this hypothesis, we evaluated glycogen, maltose, isomaltose, and trehalose (Figure 4C). Glycogen is a glucose polysaccharide with two types of bonds (α-1,4 and α-1,6) that need to be cleaved to liberate glucose for metabolism. When spores were exposed to excess glycogen as the sole carbon source, spores germinated but did so at a slower rate than in glucose. These results suggest that spores can access metabolizable carbon in glycogen but at a limiting rate, thus not allowing for fully efficient germination (82.8% germination efficiency relative to free glucose). In addition, spores in glycogen stalled after exiting the spore state and were unable to undergo further growth, implying that glycogen was not a suitable carbon source for yeast growth after germination. This was supported by the inability of yeast to grow in culture when glycogen was the sole carbon source (5.4% yeast growth relative to glucose). In the presence of maltose and isomaltose, spores germinated at rates equivalent to glucose (101.8% and 100.0% relative germination efficiency respectively), suggesting that spores harbor enzymes capable of cleaving α-1,4 and α-1,6 glycosidic bonds but that these enzymes were not accessible for the breakdown of extracellular glycogen. Yeast showed less optimal abilities to utilize maltose and isomaltose showing 78.5% and 51.8% relative yeast growth, respectively.

Another common storage sugar, trehalose, is a glucose disaccharide with a 1,1-glycosidic bond, which requires trehalase enzymes for breakdown. Spores in the presence of trehalose as the sole carbon source germinated but at a much slower rate than in glucose (46.9% relative germination efficiency). We observed that spores in trehalose take 42 hours to reach the equivalent of 12 hours of efficient germination in glucose. This spore population does not show a slow start like the one seen with raffinose; rather, germination appears to initiate immediately and progress at a constant, but slow rate. These data suggest that *Cryptococcus* spores can transport and/or hydrolyze trehalose at a slow rate from the beginning of germination. Trehalose did not support robust yeast growth as the sole carbon source, demonstrating only 3.5% relative yeast growth. Together, these results indicate that spores can break down, transport and/or metabolize external glycogen and trehalose in a spore-specific manner that is not active in yeast.

There are two known trehalase enzymes present in *Cryptococcus*: Nth1 is a cytosolic trehalase essential for yeast growth on plates containing trehalose as the sole carbon source, and Nth2 is a putatively secreted trehalase that is largely dispensable for yeast growth [26]. While *NTH1* transcript levels are comparable between yeast and spores, *NTH2* transcripts are highly abundant in spores and largely absent in yeast [26]. To assess the roles of these trehalase enzymes in germination, we evaluated wild type, *nth1*Δ, *nth2*Δ, and *nth1*Δ *nth2*Δ spores in the presence of either glucose or trehalose as the sole carbon source over 42 hours (Figure 4D). In the presence of glucose, spores from each background demonstrated equivalent rates of germination; however, in the presence of trehalose as the sole carbon source, germination was affected in both trehalase mutants. *nth1*Δ spores demonstrated a modest phenotype in which germination efficiency was roughly equivalent to wild type spores but was slightly less synchronous than wild type (showing a broader spread in sizes and shapes). In contrast, *nth2*Δ spores demonstrated a markedly reduced level of germination, plateauing with ∼35% of the population transitioned out of the spore state. The *nth1*Δ *nth2*Δ spores demonstrated a phenotype identical to that of *nth2*Δ spores. These results suggest that Nth1 is a constitutively expressed and active enzyme that hydrolyzes intracellular trehalose, whereas Nth2 is expressed in a spore-specific manner and secreted extracellularly to enable the breakdown of exogenous trehalose.

## Discussion

Spores are a critical cell type for the survival of many fungi, including the majority of human fungal pathogens [2–4]. Here we determined the molecular and metabolic requirements for germination of *Cryptococcus* spores. We identified morphological changes that occur across 12 hours of germination in glucose, showing key surface composition changes during this differentiation process. Using inhibitors of key eukaryotic processes, we identified temporal associations between molecular and morphological events and determined that carbon metabolism pathways (both glycolysis and oxidative phosphorylation) were required from the beginning of germination. These data indicated that, unlike spore germination in other organisms, bioavailable carbon is the primary requirement for the efficient induction and maintenance of germination in *Cryptococcus* spores. Furthermore, the sources of bioavailable carbon are more diverse for spores than they are for yeast, and they appear to need to be both transported into the cell and hydrolyzed to initiate and sustain germination. This demonstrates that carbon sources are required fuels for sustaining germination, rather than simply “triggers” that initiate an autonomous germination program seen in other systems.

The most well understood inducers and regulatory mechanisms of germination have been discovered in bacteria such as *B. subtilis* where L-alanine can induce germination through the binding of specific cell surface receptors [27]. This suggests that nutrients can be sufficient as signals to trigger germination without the need for exogenous metabolizable nutrients. This has also been demonstrated in the fungi *Aspergillus niger* and *Aspergillus nidulans* in which the first steps in conidial germination can be induced by non-metabolizable glucose analogs (e.g., 2-DG) [28,29]. In the case of *S. cerevisiae*, germination is successfully induced and maintained even after the removal of glucose, demonstrating a ‘commitment’ step similar to those observed in bacteria [16]. In contrast, we report here that in *Cryptococcus* non-metabolizable glucose analogs (including 2-DG) are insufficient to induce germination, suggesting that an initial metabolic priming is necessary to reach a ‘commitment’ step. Not only is 2-DG not a usable carbon source to induce germination, but due to its ability to inhibit glycolysis, it also actively inhibits germination. Similarly, when exposed to limited glucose, *Cryptococcus* spores germinate efficiently until they stall in a concentration dependent manner, suggesting that in this system, nutrient signals are not sufficient to reach a ‘commitment’ step, but rather nutrients act as a fuel for germination. As demonstrated here with glucose as the fuel, germination proceeds efficiently until the fuel is fully metabolized, germinating spores can then remain in a partially germinated state for at least 16 hours, and they resume germination when the fuel supply is replenished. While studies on the role of nutritional triggers in fungal spore germination are generally sparse and do not convey germination kinetics, it is clear that different fungal spores respond to different triggers to induce and maintain germination. In *Cryptococcus*, metabolizable carbon sources are necessary fuel for both the induction and maintenance of germination.

One of the potential benefits of this discovery is the ability to use carbon source limitation to stall spores at specific stages of germination (Figure 3). Partially germinated populations of spores could be used to evaluate the different molecular events that occur over the course of germination or to evaluate the role of partially germinated spores in pathogenesis. In addition to carbon limitation, the complex sugars characterized in this study can be used as tools to better understand the regulatory mechanisms of spore germination. Different complex sugars led to different behaviors, suggesting intrinsically different mechanisms of breakdown. For example, sucrose is readily used by spores during germination, suggesting that surface invertases or efficient sucrose transporters exist in dormant spores. This is in contrast to lactose, which is not a viable carbon source for initiating or sustaining germination, suggesting that *Cryptococcus* does not produce a β-galactosidase capable of breaking down lactose. A carbon source that elicited a unique phenotype during germination was the trisaccharide raffinose. In raffinose as the sole carbon source, spores remained morphologically ungerminated for the first 6 to 8 hours and then initiated and underwent efficient germination. One possibility for this result is that spores can detect raffinose in their environment and respond by producing the transporters and/or enzymes necessary for raffinose utilization (and this takes approximately 8 hours). Due to this unique feature, raffinose could be a powerful tool to better understand how dormant spores sense their environments. All together, the profiles of carbon source utilization by spores indicate both breadth and flexibility, supporting a robust response to nutrient acquisition as might be anticipated by an environmental fungus primed for survival in diverse environments.

One major outcome of this work was the identification of (at least 3) spore-specific metabolic pathways. Spores demonstrated a clear ability to metabolize glycogen, trehalose, and inositol as sole carbon sources, whereas yeast did not. Glycogen is readily utilized by spores during germination, indicating that *Cryptococcus* spores have the enzymes necessary to metabolize this sugar at hand; however, germinated spores (yeast) stall after germination, which is consistent with the inability of yeast to use glycogen as a carbon source during replication. This finding suggests that spores either stop producing or secreting the components required to utilize glycogen or actively inhibit the use of glycogen as fuel post-germination under these conditions. Further investigation of this process will be required to discern how and why a new cell type would halt growth and replication in the presence of a previously accessible and currently available fuel source.

In contrast, the disaccharide trehalose supported immediate germination at a modest but consistent rate until yeast growth initiation at which time vegetative growth nearly stalled and became extremely slow. It was shown previously that there are two trehalases present in *Cryptococcus* that contribute to trehalose utilization, Nth1 and Nth2. *NTH1* transcripts were present in both spores and yeast, but *NTH2* transcripts were highly abundant in spores and during sexual development and largely absent in actively replicating yeast [26]. Here we discovered that Nth2, a putative secreted trehalase, is the enzyme responsible for the ability of spores to utilize trehalose during germination, suggesting that spores can use both intracellular and extracellular reservoirs of trehalose, whereas yeast can use only intracellular stores. The use of extracellular trehalose by spores is also interesting because in contrast to the germination delay seen in raffinose, germination in trehalose is constant (albeit inefficient) from the beginning of germination. This suggests that there is no regulatory response to increase Nth2 production in the presence of available trehalose, or there are other rate-limiting steps that prevent the use of exogenous trehalose (e.g. limited Nth2 secretion or absence of trehalose import). Trehalose metabolism in spores is constitutively programmed to use exogenous trehalose during germination and not during vegetative growth. This is the first reported example of a spore-specific enzyme identified in fungi and suggests that spore germination is not simply a modified cell cycle for exiting a quiescent state, but rather, germination represents a highly adapted differentiation process that engages unique molecular pathways.

Finally, the discovery that spores readily utilize inositol during germination when yeast cannot during growth has potential implications for spores as infectious propagules. Inositol has been shown to be important in the *Cryptococcus* life cycle, which is clearly demonstrated by *Cryptococcus* species housing 10-11 inositol transporters in their genomes (as opposed to 1 or 2 like most other known fungi) [19]. Inositol has been shown to induce sexual development but has also been implicated in virulence, which is linked to the high concentration of inositol in the brain, the main site of cryptococcosis [19–21]. It has been demonstrated previously that spores readily disseminate out of the lung and that spore-mediated infections lead to significantly higher fungal burden in the brains of mice than in yeast-mediated infections [30]. The potential role of inositol in spore-specific disease kinetics are yet to be investigated, but one possibility is that this unique metabolic potential of spores could give spores an advantage in brain colonization. It is clearly beneficial for spores to have broad metabolite utilization to enable them to readily adapt to new environments to which they disperse; however, the reasons behind the limited ability of *Cryptococcus* to use an available carbon source for growth when the metabolic pathways to use it are available remain to be determined. Overall, distinguishing the metabolic potential of spores promises to identify and inform fungal-specific pathways and processes that influence survival in new environments, including the mammalian host.

## Materials and Methods

### Strains and Strain Manipulation

*Cryptococcus neoformans* serotype D (*C. deneoformans)* strains JEC20, JEC21, CHY1705, CHY1710, CHY2082, CHY2083, CHY2180 and CHY2182 were handled using standard techniques and media as described previously [26, 31, 32]. *Cryptococcus* spores were isolated from cultures as described previously [10]. Briefly, yeast of both mating types (JEC20 and JEC21, CHY1705 and CHY1710, CHY2082 and CHY2083, or CHY2180 and CHY2182) were grown on yeast-peptone-dextrose (YPD) medium for 2 days at 30°C, combined at a 1:1 ratio in 1X phosphate buffered saline (PBS), and spotted onto V8 pH 7 agar plates. Plates were incubated for 5 days at 25°C, and crosses were scraped up, resuspended in 75% Percoll in 1X PBS, and subjected to gradient centrifugation. Spores were recovered, counted using a hemocytometer, and assessed for purity by visual inspection.

### Spore Germination Microscopy

Spores were resuspended in YPD broth and incubated at 30°C with shaking at 200 RPM. Aliquots of germinating spores were removed every four hours for 12 hours and fixed immediately. Samples were fixed by incubation with 2.5% glutaraldehyde and 2% formaldehyde in PBS.

### Scanning electron microscopy (SEM)

Dormant and germinating spores were washed free of fixative, applied to poly-L-lysine-coated coverslips (Sigma), allowed to adhere for one hour, and then soaked in 4% osmium tetroxide for one hour. Coverslips were then dehydrated by submerging coverslips in a series of increasing concentrations of ethanol (EtOH) (25%, 50%, 75%, 95%, and 100% three times) allowing for 10 minutes of equilibration at room temperature between ethanol treatments. Coverslips were washed in hexamethyldisilazane (HMDS, Sigma) three times by submerging and allowing 10 minutes for equilibration between HDMS treatments. Coverslips were then air dried at room temperature, and sputter coated with palladium. A Hitachi S-570 scanning electron microscope was used to visualize the samples.

### Transmission electron microscopy (TEM)

Fixed dormant and germinating spores were post-fixed in 1% osmium tetroxide, 1% potassium ferrocyanide in 0.1M PB for 1 hour at room temperature and dehydrated in EtOH. Samples were infiltrated with increasing PolyBed 812 (Polysciences Inc.) and propylene oxide and placed in embedding capsules (BEEM) and fresh PolyBed 812 for 48 hours at 60°C. Embedded spores were sectioned on a Leica EM UC6 ultramicrotome at 90nm, and sections were post-stained in uranyl acetate and lead citrate. Samples were viewed at 80 kV on a Philips CM120 transmission electron microscope.

### Fluorescence microscopy

Fixed dormant and germinating spores were suspended in 10 mM 4-(2-hydroxyethyl)-1-piperazineethanesulfonic acid (HEPES) buffer with 0.15 M NaCl, 0.1 mM CaCl_2_ and 0.1 mM MnCl_2_. Fluorescein conjugated *Datura stramonium* then added at a 1:50 dilution, and anti-GXM antibody (F12D2) conjugated to phycoerythrin (PE) was added at a 50 µg/mL concentration. Counterstaining was with 4’-6-diamidino-2-phenylindole (DAPI) (Invitrogen) at 20 µg/mL. Cells were incubated for 20 minutes on ice and then washed twice in HEPES buffer prior to visualization on a Zeiss AxioscopeA.1 using standard filters. Single channel images were merged using Photoshop CS3.

### Spore nuclei quantification

Spores were stained with 2µM SYTOX Green and 25 µg/mL calcofluor white for 30 minutes and washed with PBS. Spores were visualized on a Zeiss AxioscopeA.1 using standard filters, and red and green fluorescent puncta were scored per spore.

### Quantitative germination assays

Germination assays were performed based on the modified Barkal et al. 2016 assay described in Ortiz et al 2021 [8,9]. Briefly, 384 well plates (Thermo Scientific: 142762) were loaded with 10^5^ spores per well. At time 0 hours, SD medium (synthetic medium + 2% D-glucose) containing the compounds of interest was added to the sample (final volume of 40 µL), unless specified otherwise. Spores were germinated at 30°C in a humidified chamber, and the same ∼5 x 10^3^ cells were monitored every 2 h for 16 h. Changes in length of experiments and alternative time intervals were performed in specific experiments, as specifically indicated. Imaging was performed on a Ti2 Nikon microscope, and each condition was visualized in a minimum of two individual wells with three fields of view acquired from each well. All images were analyzed as described previously based on cell shape and size using ImageJ. The population ratios of spores, intermediates, and yeast were determined. Error bars in plots are based on the variation among all fields of view acquired. Level of germination was determined by quantifying the decrease in the proportion of spores in a population, and rates were quantified by determining the change in this proportion over time. For all two-dimensional histograms, the x-axis represents size from 0 µm^2^ to 25 µm^2^, and the y-axis represents aspect ratios (width/length) from 0.4 to 1.

### Inhibitors and concentrations

All inhibitor compounds were tested initially at 80 µM; however, concentrations were changed on a case-by-case basis for subsequent experiments. All assays and controls were performed with a final concentration 0.8% DMSO (the compound solvent) unless specified otherwise. Antimycin A (80 µM) (Santa Cruz Biotechnology, Inc: sc-202467A), actinomycin D (160 µM) (Dot Scientific, Inc: DSA10025-0.005), CPTH2 (20 µM – 80 µM) (Fisher Scientific: AAJ65939LB0), aphidicolin (200 µM) (Fisher Scientific: NC9906396), trichostatin A (40 µM – 80 µM) (VWR: 103743-284), bortezomib (650 µM) (Fisher Scientific: 102988-732), 5-azacytidine (50 µM) (VWR: 75844-560), sodium butyrate (80 µM) (Fisher Scientific: AAA1107906), hydroxyurea (80 µM) (Fisher Scientific: 501151572), nocodazole (80 µM) (Fisher Scientific: AC358240100), paclitaxel (80 µM) (Fisher Scientific: 501445376), cytochalasin A (80 µM) (Santa Cruz Biotechnology, Inc: SC-204705), DIP (100 µM) (Fisher Scientific: AC117500100), and 1,10-phenanthroline (10 µM – 80 µM) (Sigma-Aldrich: 131377-5G) were all tested in 0.8% DMSO. Pladienolide B (20 µM – 80 µM) (Santa Cruz Biotechnology, Inc: SC-391691) was tested in 1.6% DMSO. 2-DG (125 mM – 500 mM) (Sigma-Aldrich: D8375-1G) and tetrathiomolybdate (TTM) (50 µM) (VWR: 103464-038) were tested with no DMSO (water soluble).

### Glucose limitation

Spores were germinated and assessed using the quantitative germination assay for 20 hours in the presence of D-glucose (concentrations from 1.7 µM to 111mM) in S medium. After 20 hours, 111mM D-glucose was added to each condition (40 µL) and germination was assessed for an additional 16 hours or until spores were fully germinated.

### Carbon source testing

All carbon sources were tested at 111mM or equivalent of 111mM of glucose subunits for non-numerical polymers (glycogen and cellulose) in synthetic medium without supplemented glucose (S medium). Relative percent spore germination was determined based on 16 hours of germination relative to the 111mM glucose control.

### Yeast replication experiments

JEC20 and JEC21 strains were grown to stationary phase in liquid YPD culture overnight at 30°C with shaking then washed with 1 X PBS. Resuspended yeast were mixed in a 1:1 ratio and diluted to an OD_600_ of 0.2 in Synthetic medium (S) with carbon sources at a concentration of 111mM in 96-well plates. Growth was monitored at 30°C with shaking by OD_600_ measurements every three minutes for twenty-four hours on a Tegan Infinite 200 Pro instrument (#30052730). The growth curves were then plotted, and percent growth was calculated based on final OD_600_ values. Relative percent growth was determined at the 16-hour time point as compared to glucose control.

## Supplemental Figures

**Figure S1.**
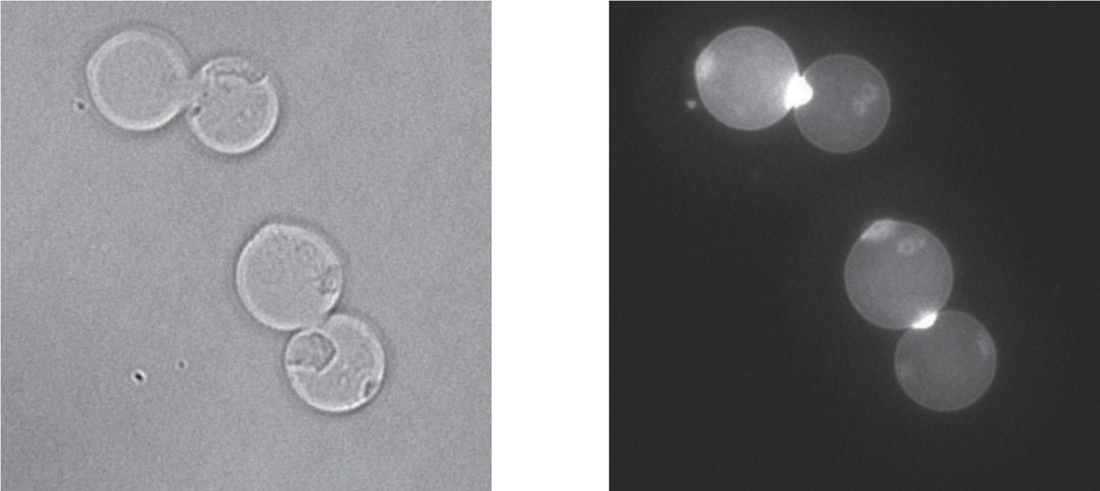
Bud scar staining of *Cryptococcus* yeast demonstrates bipolar budding. Left: DIC image of actively budding yeast at 1000X magnification. Right: Fluorescence microscopy image of same yeast showing high intensity of calcofluor white staining at bud scars.

**Figure S2.**
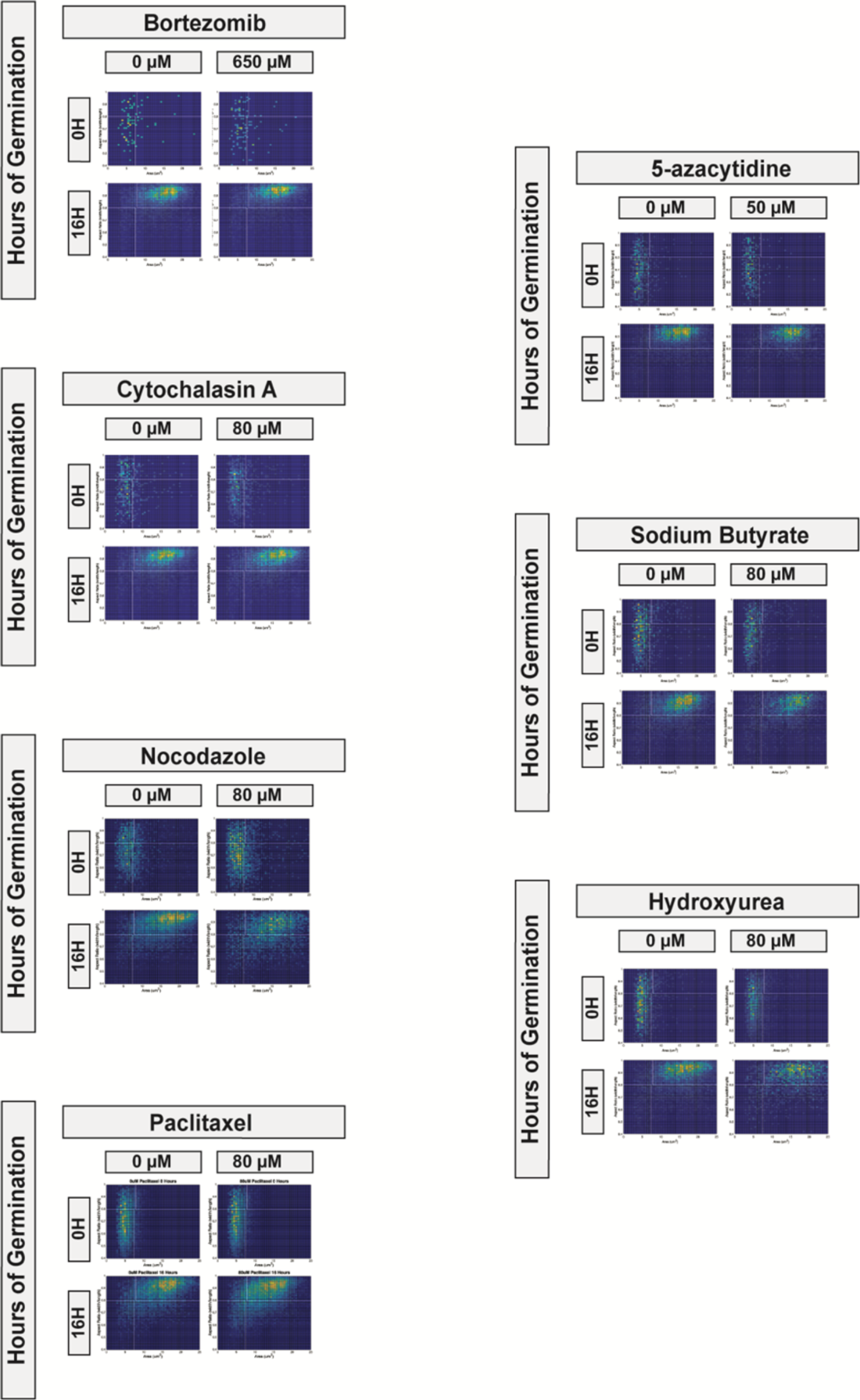
A variety of inhibitors of eukaryotic processes did not inhibit germination. Germination profiles at 0 and 16 hours of spores germinating in the presence of bortezomib (650 μM), 5-azacytidine (50 μM), sodium butyrate (80 μM), hydroxyurea (80 μM), nocodazole (80 μM), paclitaxel (80 μM), or cytochalasin A (80μM). For all two-dimensional histograms, the x-axis represents size from 0 μm^2^ to 25 μm^2^, and the y-axis represents aspect ratio from 0.4 to 1.

**Figure S3.**
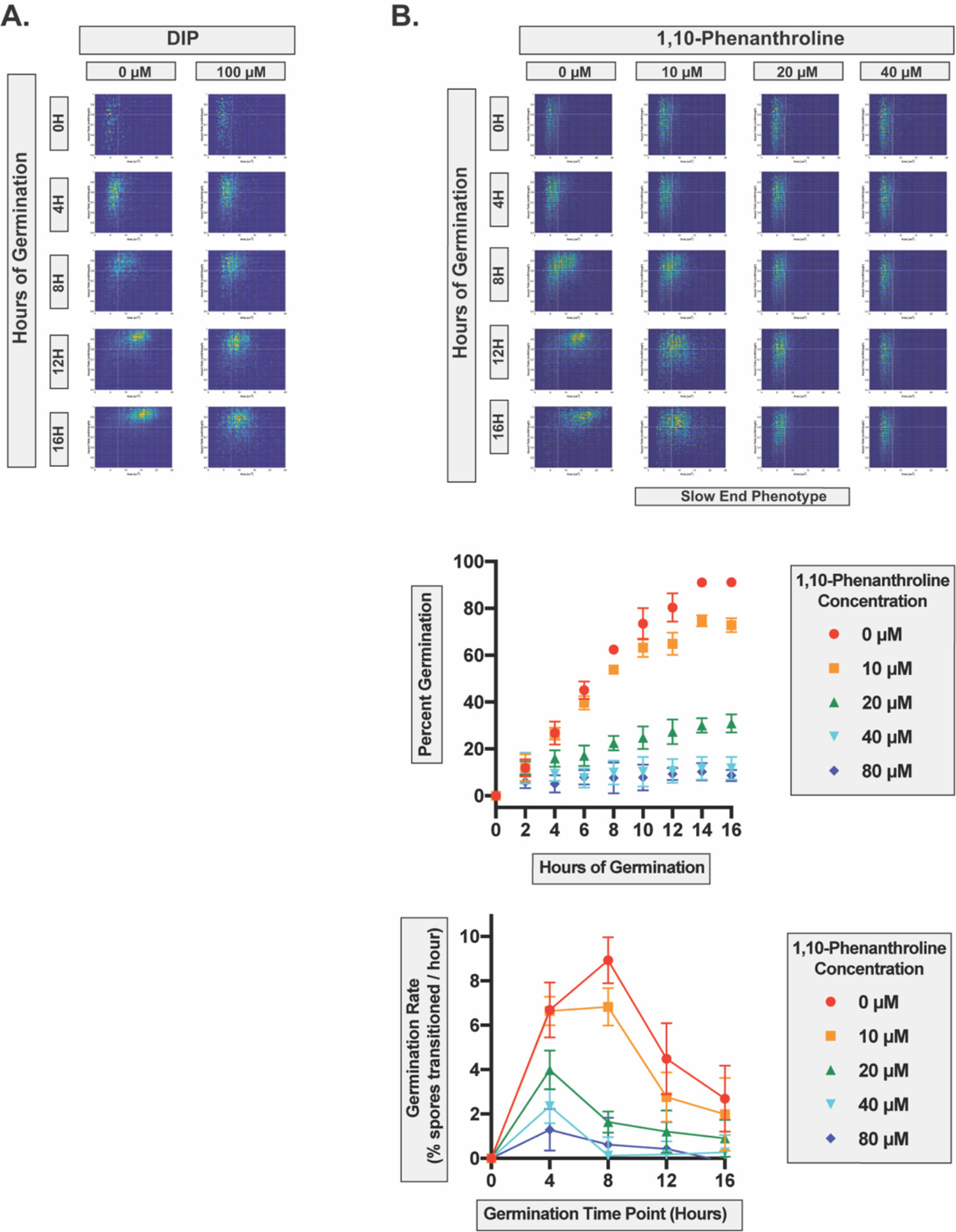
The non-specific metal chelators DIP and 1,10-phenanthroline induce a “slow end” phenotype. A) Germination profiles of spores at 0, 4, 8, 12 and 16 hours germinating in the presence of 100 μM DIP. B) Germination profiles of spores at 0, 4, 8, 12 and 16 hours germinating in the presence of 10, 20 or 40 μM of 1,10-phenanthroline, along with quantitative assessments of germination and germination rates (for 10, 20, 40 and 80 μM 1,10-phenanthroline). For all two dimensional histograms, the x-axis represents size from 0 μm^2^ to 25 μm^2^, and the y-axis represents aspect ratio from 0.4 to 1.

**Figure S4.**
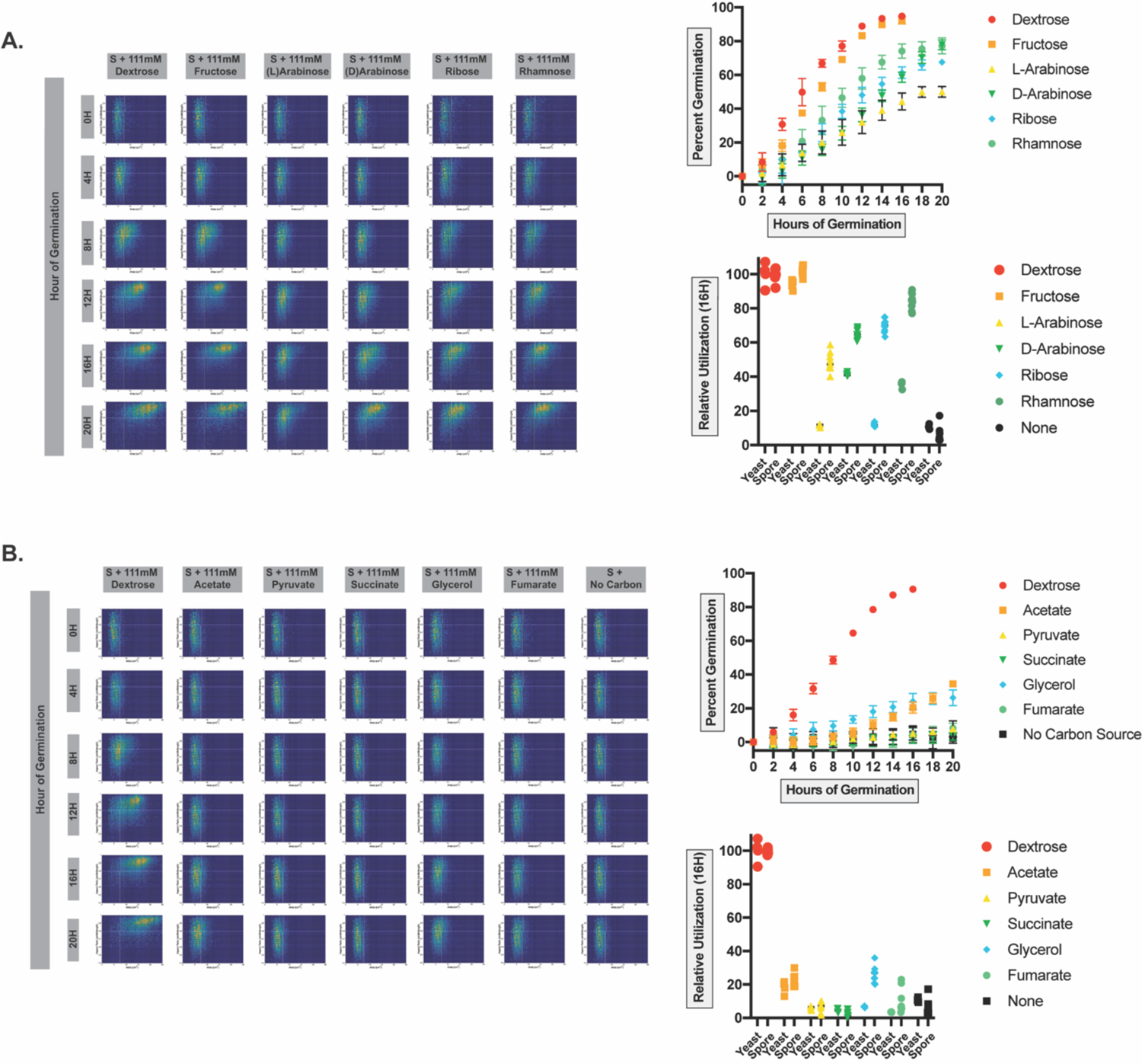
Spores and yeast demonstrate different metabolic potentials in a variety of carbon sources. **A)** Germination profiles of spores at 0, 4, 8, 12, 16 and 20 hours of germination in the presence of a variety of simple sugars (111mM) as sole carbon sources including fructose, L-arabinose, D-arabinose, ribose and rhamnose along with the quantitative assessments of germination and relative nutrient utilization for spore germination and yeast replication under these conditions (relative to dextrose). **B)** Germination profiles of spores at 0, 4, 8, 12, 16 and 20 hours of germination in the presence of a variety of non-sugar carbon sources (111mM) as sole carbon sources including acetate, pyruvate, succinate, glycerol, and fumarate along with the quantitative assessments of germination and relative nutrient utilization for spore germination and yeast replication under these conditions (relative to dextrose). For all two-dimensional histograms, the x-axis represents size from 0 μm^2^ to 25 μm^2^, and the y-axis represents aspect ratio from 0.4 to 1.

**Figure S5.**
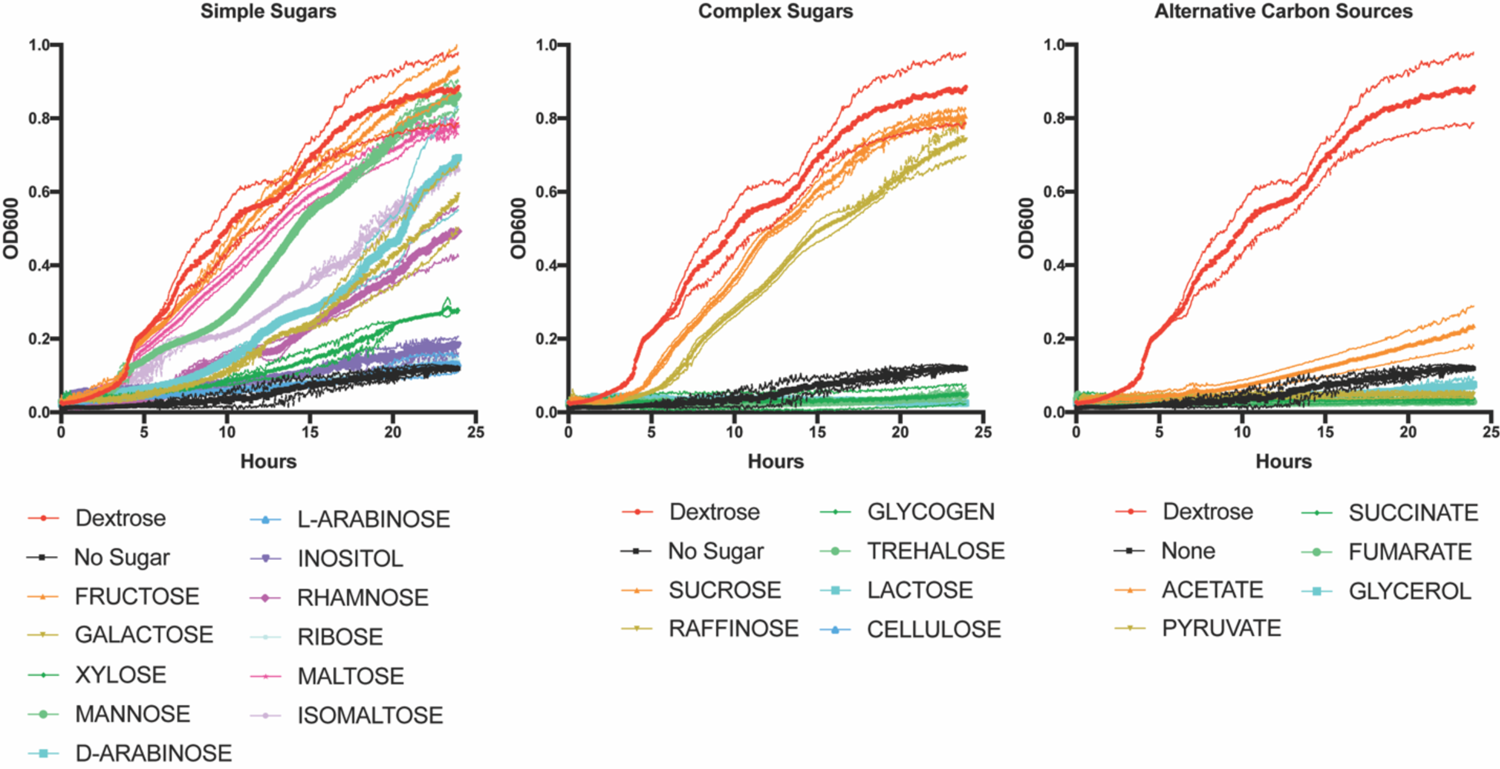
The efficiency of *Cryptococcus* yeast replication varies widely across different carbon sources. Optical density measurements of yeast (JEC20 and JEC21) replicating in the presence of indicated carbon sources (111mM) for 24 hours.

---

**Table S1.** Summary of relative carbon source use during germination and growth. Germination and growth were tested in 21 carbon sources and compared to glucose. Efficiency of spore germination was defined as the percentage of spores escaping the spore state after 16 hours in germination conditions at 30°C. Percent utilization is the germination efficiency of a C source relative to that of glucose (Germination Efficiency of test C/Germination Efficiency of glucose). Yeast growth in each carbon source was measured by growing liquid cultures in rich medium (YP+carbon source) and taking OD600 readings after 16 hours at 30°C. Percent utilization is the OD600 of test C source/OD600 of glucose. Growth in glucose was set to represent 100% efficiency.

| Carbon Source | % Utilization (relative to glucose) |  |
| --- | --- | --- |
|  | Spore Germination | Yeast Growth |
| Glucose | 100 | 100 |
| Fructose | 102.0 | 93.9 |
| Maltose | 101.8 | 83.8 |
| Sucrose | 101.1 | 86.6 |
| Isomaltose | 100.0 | 51.8 |
| Mannose | 90.4 | 78.5 |
| Rhamnose | 84.4 | 35.4 |
| Inositol (myo) | 83.0 | 16.0 |
| Glycogen | 82.8 | 5.4 |
| Ribose | 69.9 | 12.0 |
| D-Arabinose | 64.7 | 41.3 |
| Galactose | 55.7 | 36.1 |
| Raffinose | 54.3 | 70.2 |
| Xylose | 50.4 | 21.2 |
| L-Arabinose | 48.9 | 11.5 |
| Trehalose | 46.9 | 3.5 |
| Glycerol | 26.2 | 6.7 |
| Acetate | 22.5 | 18.2 |
| Lactose | 18.4 | 4.3 |
| Fumarate | 9.2 | 3.5 |
| Pyruvate | 5.6 | 6.1 |
| Succinate | 1.8 | 4.5 |
| Cellulose | 0 | 0 |

